# ‘Pscore’ - A Novel Percentile-Based Metric to Accurately Assess Individual Deviations in Non-Gaussian Distributions of Quantitative MRI Metrics

**DOI:** 10.1101/2023.12.10.571016

**Authors:** Rakibul Hafiz, M. Okan Irfanoglu, Amritha Nayak, Carlo Pierpaoli

## Abstract

**BACKGROUND:** Quantitative MRI metrics could be used in personalized medicine to assess individuals against normative distributions. Conventional Zscore analysis is inadequate in the presence of non-Gaussian distributions. Therefore, if quantitative MRI metrics deviate from normality, an alternative is needed.

**PURPOSE:** To confirm non-Gaussianity of diffusion MRI (dMRI) metrics on a publicly available dataset, and to propose a novel percentile-based method, ‘Pscore’ to address this issue.

**STUDY TYPE:** Retrospective cohort

**POPULATION:** 961 healthy young-adults (age:22-35 years, Females:53%) from the Human Connectome Project

**FIELD STRENGTH/SEQUENCE:** 3-T, spin-echo diffusion echo-planar imaging, T1-weighted: MPRAGE

**ASSESSMENT:** The dMRI data were preprocessed using the TORTOISE pipeline. Forty-eight regions of interest (ROIs) from the JHU-atlas were redrawn on a study-specific diffusion tensor (DT) template and average values were computed from various DT and mean apparent propagator (MAP) metrics. For each ROI, percentile ranks across participants were computed to generate ‘Pscores’– which normalized the difference between the median and a participant’s value with the corresponding difference between the median and the 5^th^/95^th^ percentile values.

**STATISTICAL TESTS:** ROI-wise distributions were assessed using Log transformations, Zscore, and the ‘Pscore’ methods. The percentages of extreme values above-95^th^ and below-5^th^ percentile boundaries (PEV_>95_(%),PEV_<5_(%)) were also assessed in the overall white matter. Bootstrapping was performed to test the reliability of Pscores in small samples (n=100) using 100 iterations.

**RESULTS:** The dMRI metric distributions were systematically non-Gaussian, including positively skewed (e.g., mean and radial diffusivity) and negatively skewed (e.g., fractional and propagator anisotropy) metrics. This resulted in unbalanced tails in Zscore distributions (PEV_>95_≠5%,PEV_<5_≠5%) whereas ‘Pscore’ distributions were symmetric and balanced (PEV_>95_=PEV_<5_=5%); even for small bootstrapped samples (average 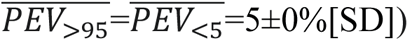.

**DATA CONCLUSION:** The inherent skewness observed for dMRI metrics may preclude the use of conventional Zscore analysis. The proposed ‘Pscore’ method may help estimating individual deviations more accurately in skewed normative data, even from small datasets.

## INTRODUCTION

Neuroimaging studies in clinical research typically rely on group-level analyses, delineating summary outcomes that differentiate a patient cohort from a group of healthy controls. However, in clinical practice, there is a need to assess individual patients. This is typically done by building a model to evaluate the individual subject against a normative sample (1–3). Quantitative MRI metrics can provide normative data against which individual deviations can be assessed. Therefore, once the accuracy and reliability are verified, quantitative MRI metrics could become highly relevant for clinical assessment of individual patients.

Individual assessments against a normative distribution are often performed using Zscores in clinical and neuroimaging paradigms (2–4). However, a thorough assessment of the distribution that generates the normative dataset itself is rarely performed. An example is the Gaussianity and homogeneity of variance, a fundamental assumption for parametric models. Deviation from these assumptions can result in biased statistical inferences, for example in neuropsychological test scores (5). Quantile regressions have been adopted to mitigate this by generating normative distributions at specific percentiles (6). These can offer a solution based on distributions at selective percentile/quantile ranks and might avoid the use of conventional mean regression models that would be biased for asymmetric distributions.

The problem is pervasive even for quantitative MRI metrics. In a pilot study (n = 48), we have recently shown prominent deviation from normality and heavy-tailed distributions of several diffusion tensor (DT) and mean apparent propagator (MAP) metrics (7–10). For example, mean diffusivity (MD) had a positively skewed distribution, while fractional anisotropy (FA) and propagator anisotropy (PA) showed a negatively skewed distribution (10). In our small sample, these non-Gaussian features were inherently related to the diffusion characteristics of water in the brain, and not originating from heterogeneity in the underlying population as it has been previously reported in large scale-public datasets (1,3). For example, for the UK-Biobank data, Fraza et al. fit a warped Gaussian model to compensate for the skewness and kurtosis in the normative data before using Zscores to assess heterogeneous individuals (1,11). Another approach popularly used to address skewness is log transformation, but it is known to worsen the issue of skewness and can lead to inaccurate and biased inferences (12,13). Our initial assessment showed that even when comparing individuals within the normative sample, Zscores may show an imbalance in extreme values at the tails: with more negative extreme values for negatively skewed distributions and, on the other hand, more positive extreme values for positively skewed distributions (10). Therefore, comparing patients against such distributions using Zscores might wrongly place some of them in the extremities of the distribution and could introduce false-positive findings.

In this study, we aimed to test if the non-Gaussianity in diffusion MRI (dMRI) metrics can be reproduced in a large sample from a homogeneous group of healthy young subjects; and assess whether a percentile-based ‘Pscore’ could accurately estimate an individual’s deviation from the central tendencies of a normative distribution; and compare its accuracy against two popular normalization methods: ‘Zscore’ and ‘Log’-transformation.

## MATERIALS AND METHODS

### Participants

We used the Human Connectome Project Young Adult (HCP-YA) cohort from the S1200 series, which has at least 1113 participants with 3T MRI data available from a pool of 1200 young adults (age range: 22-35 years); who were all recruited with written informed consent following the guidelines of the institutional review board (IRB) (14). The data include structural MRI, functional (fMRI), and dMRI data along with behavioral and genetic testing. For this study, our primary interest was the dMRI data. About 100 (out of 1200) participants either did not have dMRI data or the parameter information on the dMRI acquisition was incomplete. From the remaining pool of 1100 subjects, 129 participants were excluded because the distortions in their subject-space dMRI data were beyond the scope of correction based on visual inspection. Ten out of the remaining 971 participants were ‘36+’ years of age, and they were excluded to keep the sample more homogeneous, within the 22-35 years range. Thus, the effective sample included neuroimaging data from 961 participants (53% females). All participants were scanned on the same equipment, using the same protocol.

### Diffusion MRI Protocol

The Connectome 3T Skyra scanner (Siemens Healthineers, Erlangen, Germany) was used to acquire dMRI data across 6 runs in a full session. Each run was approximately 9 minutes and 50 seconds long, with three different gradient tables, and each table was acquired with two phase-encoding directions – right-to-left and left-to-right. Each gradient table consisted of six b = 0 s/mm^2^ acquisitions and approximately 90 diffusion weighting directions. The diffusion weighting was done across three b-shells – 1000, 2000, and 3000 s/mm^2^, each with an equal number of acquisitions per run. The imaging parameters were as follows: spin-echo echo-planar imaging (EPI) sequence with repetition time (TR) = 5520 ms, echo time (TE) = 89.5 ms, flip angle = 78°, refocusing flip angle = 160°, field of view (FOV) = 210×180, matrix = 168×144, slice thickness = 1.25 mm, 111 slices acquired at 1.25 mm isotropic resolution, a multiband factor of 3, and echo spacing = 0.78ms. The T1-weighted imaging sequence included – 3D MPRAGE images acquired over a period of 7 minutes and 40 seconds at a TR = 2400 ms, TE = 2.14 ms, inversion time (TI) = 1000 ms, flip angle = 8° and FOV = 224×224 at an isotropic voxelwise resolution of 0.7 mm.

### Preprocessing

We used the TORTOISEV3 (version 3, www.tortoisedti.org) pipeline to process the dMRI data, because Irfanoglu et al. had shown considerable improvement in the dMRI metrics using this pipeline compared to the released version of the HCP dataset (15,16). In the following, we briefly describe the different stages of pre-processing: *(a) Denoising* was performed using a model-free noise mapping technique proposed by Veraart et al, with a kernel radius of 3 (17). *(b) Gibbs-ringing correction* was performed using the subvoxel-shift method, which showed improvements even without introducing additional imperfections (18). For *(c) Inter-Volume Motion and Eddy Current Correction,* we first applied a MAP-based model, as it is independent of shelled data and accurately extrapolates the unseen q-vector signals (16,19). All diffusion-weighted images (DWIs) were aligned to the ideal b = 0 s/mm^2^ image. As a final step, step *(c)* was run once more, but in this instance, the synthesized and the slice-transformed real images were used. These steps were iterated over until convergence was reached. *(d) Susceptibility-induced distortions* were considered as well, given that acquiring data at high resolution (like for the HCP) can cause severe EPI distortions (14,20). We applied the DRBUDDI approach, which is a blip-up and blip-down distortion correction technique with excellent performance (20,21). Besides using the b = 0 s/mm^2^ image, DRBUDDI also incorporates DTs and relies on an undistorted T2-weighted (T2W) structural image (20). However, the T2W images from the HCP data were not compatible with DRBUDDI; therefore, we used a machine learning-based technique called SynB0-DisCo to generate a structural image that fit well into the DRBUDDI paradigm (22). *(e) Gradient nonlinearity correction* was used as the HCP data come with “gradwarped” DWIs and gradient-deviation tensor images (14). Effects of inter-volume motion were not considered when a single gradient-deviation tensor was used for all DWIs. Therefore, we also computed the voxelwise B matrices, to take these effects into consideration (23). *(f) Signal drift correction*, given that the scan time was long, signal drift was observed in the HCP data, and therefore had to be corrected using a method proposed earlier (24). *(g) Normalization and template generation* was performed on the processed data at both the HCP isotropic resolution of 1.25 mm and at the 1 mm resolution of the processed T1-weighted (T1W) image. A DT-based registration was applied, and an atlas was generated using the 1 mm resolution data; and the DWIs were warped on to the template space using non-linear transformation (25,26).

### Diffusion MRI Metrics

We generated voxelwise maps for four DT metrics – FA, MD, axial diffusivity (AD), and radial diffusivity (RD) (27). We also generated voxelwise maps for five MAP metrics – PA, return to axis probability (RTAP), return to origin probability (RTOP), return to plane probability (RTPP), and non-Gaussianity (NG) (9). Each of these metrics carries useful quantitative information about the various diffusion behaviors of water in the brain.

### Quality Control Assessment

Quality Control assessments are vital for neuroimaging applications, especially those that involve complex interpolations and multiple preprocessing steps. We took several steps in assessing each dMRI metric map for each subject. All dMRI maps registered to the study template were first visually inspected. The maps from 961 subjects were checked for misregistration and abnormal warping by generating an in-house custom-built script using modules from SPM12 (Update Revision Number: 7771, http://www.fil.ion.ucl.ac.uk/spm/) within a MATLAB environment (Version: R2022b, MathWorks Inc., Massachusetts, USA). The script generates contour lines of all maps over the corresponding subject’s T2W image in the template space. Subjects that failed the quality control assessment were checked again and appropriate steps were taken to correct the registration issues. The datasets that could not be salvaged despite this extensive correction pipeline were removed from further analysis, leaving an effective sample of 960 and 912 participants for the DT and MAP metrics, respectively.

### Regions of Interest

To reduce the number of tests and focus on specific white matter (WM) regions, we used a set of regions of interest (ROIs) for the current study. The ROIs were inspired by the John Hopkins University (JHU) WM ROIs; however, they were manually redrawn (by A.N., over 13 years of experience in the field of neuroanatomy and neuroimaging) on an average DT brain template built from the HCP dataset to avoid issues with left/right structural asymmetry that have been reported for the original JHU ROIs (19,25,28–30). Moreover, the JHU ROIs were defined in a scalar map and used scalar-based registration, which tend to have misregistrations (25). We used a tensor-based registration, which provides better alignment (25). The JHU ROIs are quite large and therefore, careful steps were taken when the ROIs were redrawn to ensure they ‘were within’ the tracts/structures to reduce partial volume effects. The ROI labels were created and the average DT and MAP metrics were computed for each subject across all ROIs using ITK-SNAP (version 3.6.0, www.itksnap.org) (29). Figure 1 shows the ROIs in a detailed montage across all three orthogonal planes for better visualization and assessment. Table S1 in the supplement provides more details on these ROIs.

**Figure 1.**
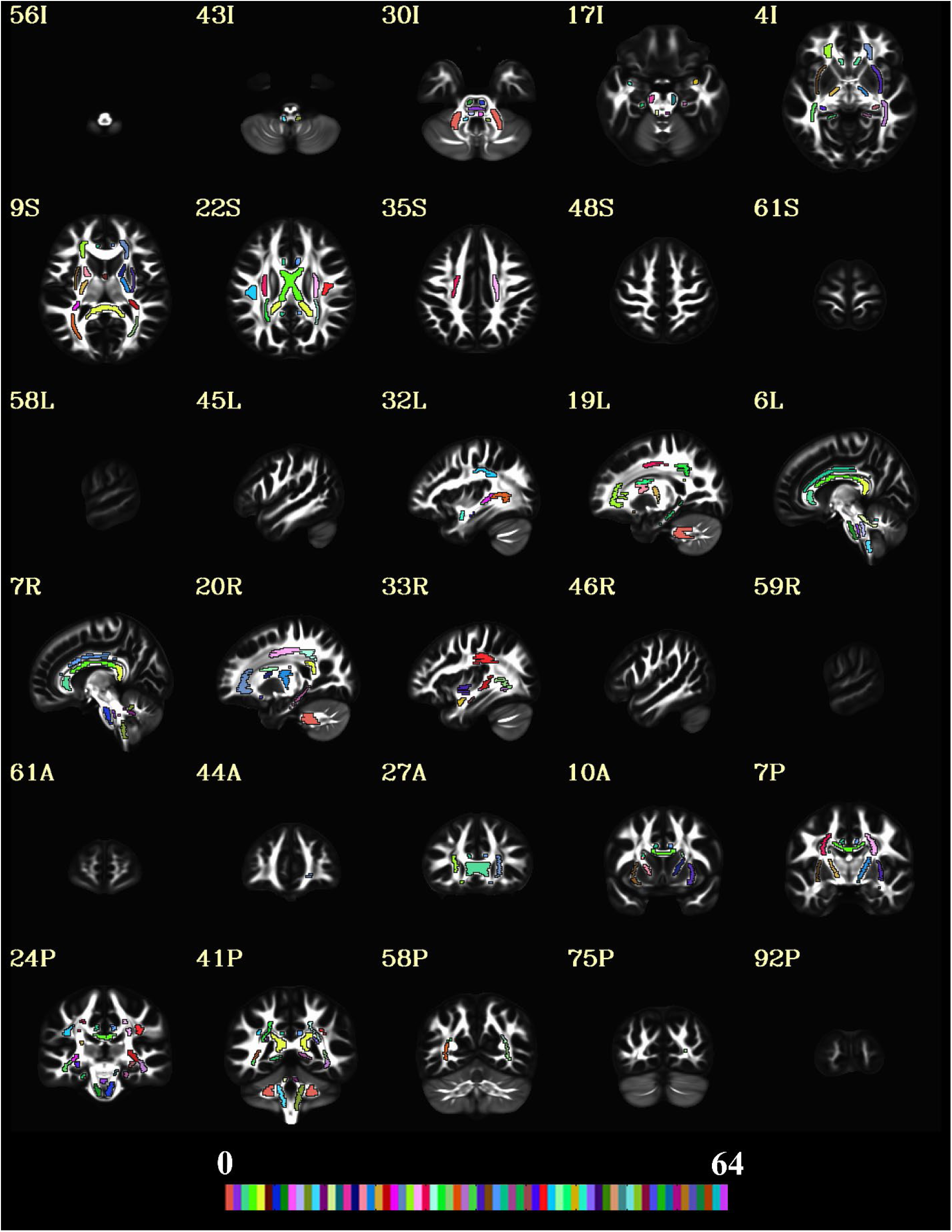
The regions of interest (ROIs) used in the current study. A 5 x 2 montage was created in all orthogonal planes to show the spatial extent of the ROIs overlaid on the average connectome diffusion tensor (FA) template. The 48 ROIs are shown using a colormap with a range of 64 colors. The slice labels with the correct direction are provided on the top-left of each image in the montage. The “A”, “P”, “I”, “S”, “L” and “R”, labels represent the anterior, posterior, inferior, superior, left and right directions, respectively.

### The Pscore Method

The first step in computing Pscores was to generate percentile ranks for each participant. They were computed within each ROI using the following formula:

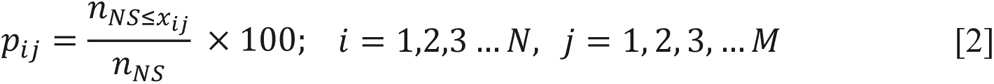

where *x*_*ij*_ represents the average dMRI metric value of the *i^th^* individual for the *j^th^* ROI, and *i* represents *1*…*N*(*960*) participants, and *j* represents *1…M*(*48*) ROIs, respectively. Furthermore, *n*_*NS*≤*x_ij_*_ represents the number of participants within the normative sample having a value ≤ *x*_*ij*_. The denominator *n*_*NS*_ represents the total number of participants in the normative sample.

After the percentile computation, each participant’s position in the ROI-wise distribution was considered to assess which side of the tail they were represented in. This was identified by the difference between the participant’s metric value and the median of the distribution. This difference was then normalized with either the difference between the median and the 5^th^ or the 95^th^ percentile edge value, depending on the participant’s position. The following equations depict how a Pscore was computed on either side of the median:

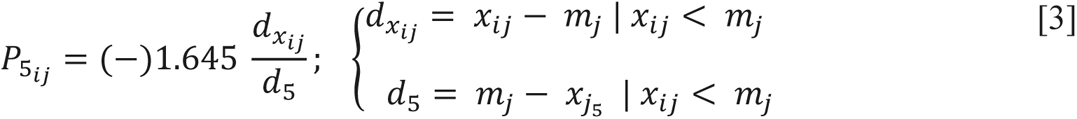

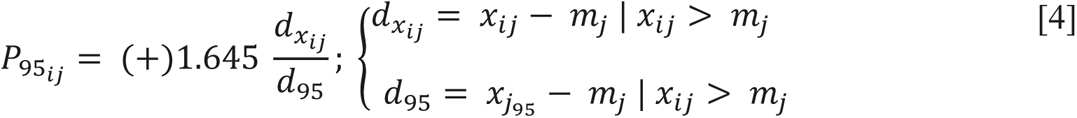

where, *d*_*x_ij_*_ is the difference between *x*_*ij*_ and the median of the distribution, *m_j_* for the *j^th^* ROI. If the metric value of the *i^th^* participant, *x*_*ij*_ < *m_j_*, then *d*_*x_ij_*_ < 0, which means that the participant was located on the left-hand tail between *m_j_* and the 5^th^ percentile edge value, *x*_*j*_5__ of the distribution. The corresponding denominator *d*_5_ was computed as the difference between *m_j_* and *x*_*j*_5__. On the other hand, *d*_*x_ij_*_ > 0, when *x*_*ij*_ > *m_j_*. This indicated that the participant’s position was on the right-hand tail between *m_j_* and the 95^th^ percentile edge value, *x*_*j*_95__ and the denominator *d*_95_ was computed as the difference between *x*_*j*_95__ and *m_j_*. Both *P*_5_*ij*__ and *P*_95_*ij*__ takes on the polarity of *d*_*x_ij_*_, generating the negative and positive scores, respectively. The ratios of these differences were then scaled by |1.645| representing the Zscore value corresponding to the 5^th^ and 95^th^ percentiles of a normal distribution. This was done to bring the Pscores to the scale of Zscores and make them comparable.

### Statistical Analysis

We analyzed the data ROI-wise in R (R Core Team (2023); Version: 4.3.2; R: A Language and Environment for Statistical Computing; R Foundation for Statistical Computing, Vienna, Austria; https://www.R-project.org/) using three normalization techniques – log transformation, standardized Zscore, and the ‘Pscore’ method proposed in this study. Histogram distributions of the raw data were used as a reference to compare the distributions from these three methods. For each histogram plot, the mean and median lines were added to assess deviations from the mode and misalignment of these moments. A normal density curve was fit on top of the histograms to assess which normalization method was closer to a Gaussian distribution. This led to four figures per ROI for each dMRI metric, generating 1728 figures (4 figures x 48 ROIs x 9 dMRI metrics).

To summarize and simplify, we took the Zscore and Pscore values from the entire sample across all ROIs and decomposed them into two single column vectors. For example, for a DT metric such as FA, each ROI had Zscores from 960 healthy individuals. This would create a vector of 46080 Zscores (960 Zscores x 48 ROIs), and similar observations were true for Pscores. For the MAP metrics, since data from 912 individuals survived the quality control step, the vector had 43776 Zscores (912 Zscores x 48 ROIs). These vectors were then used to generate two distribution plots – one for Zscores and the other for Pscores. The process was repeated for each DT and MAP metric. The log transformation method was not included in this step, as we already mentioned the issues with this approach, and it performed very poorly at the ROI level (Figure 2 and Figures S1, S2). Additionally, log-transformed values were not on the same scale as Zscores and Pscores, making them incompatible for this comparison. Since the Pscores were in the same scale as Zscores, they can be compared and assessed together. This was to showcase which of these two normalization techniques showed a statistical imbalance in the data spread across the entire WM and over the entire sample.

**Figure 2.**
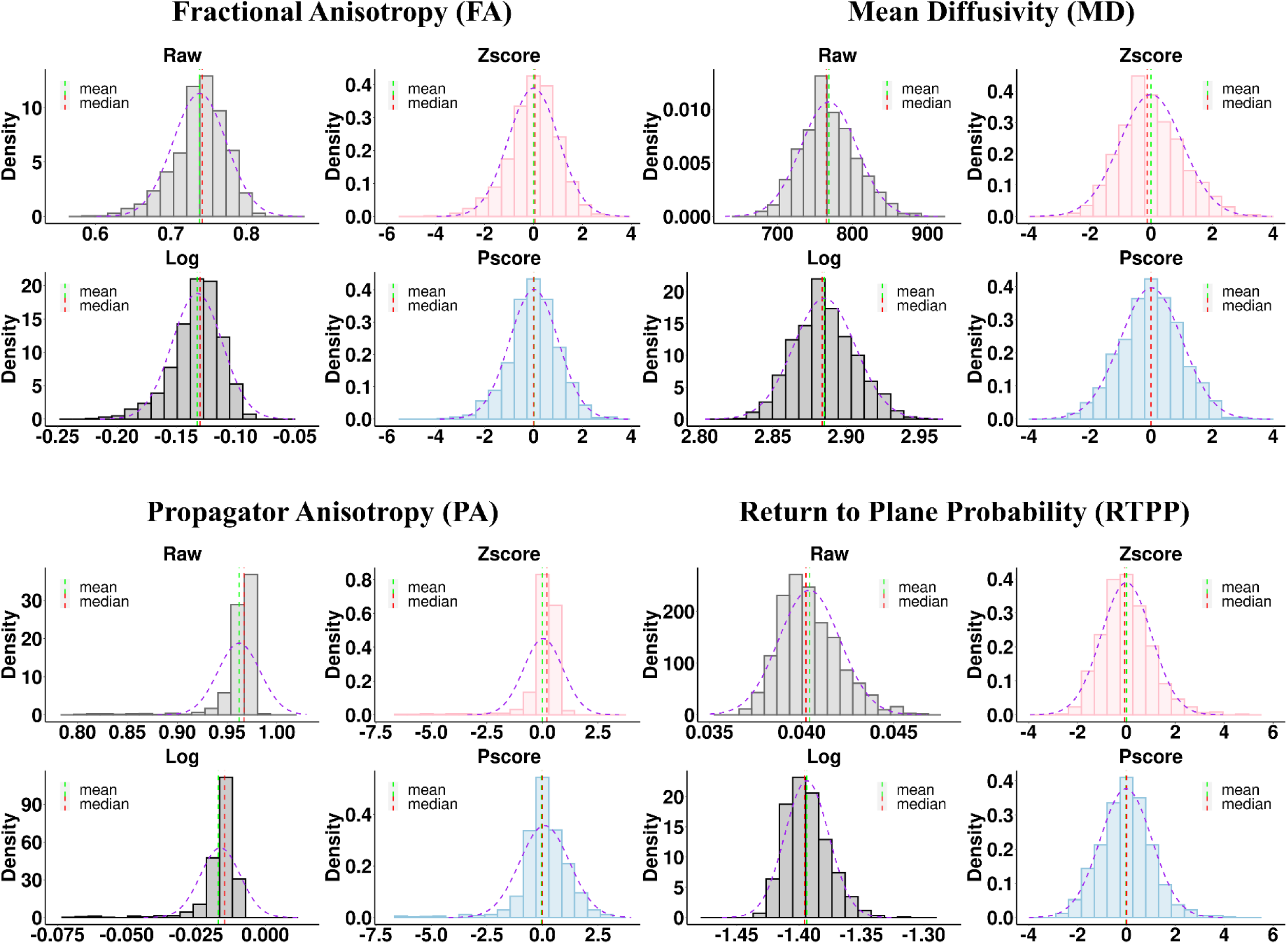
Comparing distributions across different normalization methods for two DT and two MAP metrics from a representative ROI: Body of the Corpus Callosum. The DT metrics are shown in the top row: FA (left) and MD (right); and the MAP metrics are on the bottom row: PA (left) and RTPP (right). For each metric, four distributions are shown. The ‘Raw’ (light gray) distribution panel represents the average metric values. The ‘Log’ (dark gray), ‘Zscore’ (light red) and ‘Pscore’ (light blue) distribution panels represent the logarithmic, standardized Z values and the proposed Pscores, respectively. The ‘raw’ (light gray) distributions demonstrated the presence of skew in the dMRI metrics which caused the mean, mode and median to misalign. For instance, FA (top left) and PA (bottom left) were negatively skewed; however, PA was more heavy-tailed and showed greater skewness. The mean (dashed green) appeared separated from the median (dashed red) and mode (tallest bar). Comparing the histograms with a fitted normal density curve (dashed purple) also illustrated the deviation from a Gaussian distribution. These patterns were also clearly visible for the log transformed (dark gray) and ‘Zscore’ (light red) distributions at varying levels. For both FA and PA, the ‘mean’ underestimated the most common values and appeared before the median and the mode of the distribution for ‘Raw’, ‘Log’ and ‘Zscore’ panels. However, the ‘Pscore’ panel shows that all three central tendencies coincided well and attained a closer fit to the normal density curve (dashed purple). On the other hand, MD (top right) and RTPP (bottom right) were positively skewed and the ‘mean’ overestimated the most common values for ‘Raw’, ‘Log’ and ‘Zscore’ panels; but the ‘Pscore’ panel shows the consistent alignment of the mean, mode and median and a closer fit to the normal density curve.

More importantly, for each metric, this step also helped to particularly quantify the imbalance in the number and percentage of extreme values present in the tails of these distributions. A normal Z-distribution would have 5% of extreme values above the 95^th^ percentile (Z = 1.645) and below the 5^th^ percentile (Z = −1.645) boundaries. We quantified and tabulated the number and percentage of these extreme values for Zscores and Pscores for all dMRI metrics. In the presence of a non-Gaussian distribution, the balance of 5% would be altered in the tails. This helped showcase which of these normalization methods maintained a systematic imbalance of extreme values that may lead to inaccurate assessment of individuals whose values were in the tails of the distribution.

To test how Pscores performed on smaller samples, we performed bootstrapping on the pool of HCP participants (n = 960). We ran 100 iterations, each time randomly selecting 100 participants and repeating the Zscore vs. Pscore comparison at the overall WM level. We also performed 20 iterations, independently on the dMRI metric that showed the strongest imbalance in extreme values across all WM ROIs. This was done for brevity and showcasing the reliability of the Pscore approach on the metric least expected to conform to Gaussianity.

To highlight the level of the imbalance in extreme values in Zscores compared to Pscores, we generated heatmaps, similar to the ones shown in our pilot study (10). It is an intuitive way to assess individuals through a visual representation of the level and direction in which the extreme values tend to increase. The heatmaps were generated across all ROIs for the entire population, as well as for a single iteration of the random sampling consisting of 100 participants to reduce the complexity of showcasing 92160 observations (46080 Zscores + 46080 Pscores). Since the pattern was consistent even in small samples, it can be shown with much better clarity for 100 participants with 9600 observations (4800 Zscores + 4800 Pscores).

## RESULTS

### ROI-wise Comparison of Distributions Across Different Methods

Figure 2 shows some examples of distributions from the normalization techniques tested, within the body of the Corpus Callosum (BCC). Distributions from only four dMRI metrics are shown for one ROI; however, assessments were made across each ROI per dMRI metric. The FA and PA values showed a negative skew, while MD and RTPP showed a positive skew. On the contrary, for all Pscore distributions, the mean, mode, and median appeared well aligned and they tended to fit closely to the normal density curve. Therefore, compared to the other normalization methods tested, and particularly the standardized Zscore approach, Pscores provided a more symmetric distribution per ROI. The supplementary material provides more examples from other dMRI metrics across other ROIs (Figures S1, S2).

### Zscore versus Pscore Distributions Across All ROIs

Figure 3 shows the Zscore and Pscore distributions for each dMRI metric, generated from the data across all WM ROIs comprising the overall WM. In general, Zscores showed an overall imbalance of positive and negative values for all metrics (less prominent for AD). Zscore distributions for FA, PA and NG showed an overall negative skew with < 50% negative and > 50% positive values. This pattern was observed to be progressively worsening for NG with 54% and for PA with 59% positive values, respectively. However, Pscores from all the three metrics maintained a balanced number of 50% negative and positive values. The rest of the DT and MAP metrics showed the opposite trend for Zscores, with > 50% of total observations being negative. For example, both MD and RTAP had 51% of all the observations as negative values. Therefore, MD had at least 460 (1% of 46000) and RTAP had at least 437 (1% of 43776) more negative Zscores than positive Zscores in their respective distributions. It was worse for RD, RTOP, and RTPP, because each of their distributions had 52% negative values, indicating twice as many negative values than MD and RTAP. All Pscore distributions, on the other hand, maintained a balanced distribution of 50% positive and negative values for every single dMRI metric.

**Figure 3.**
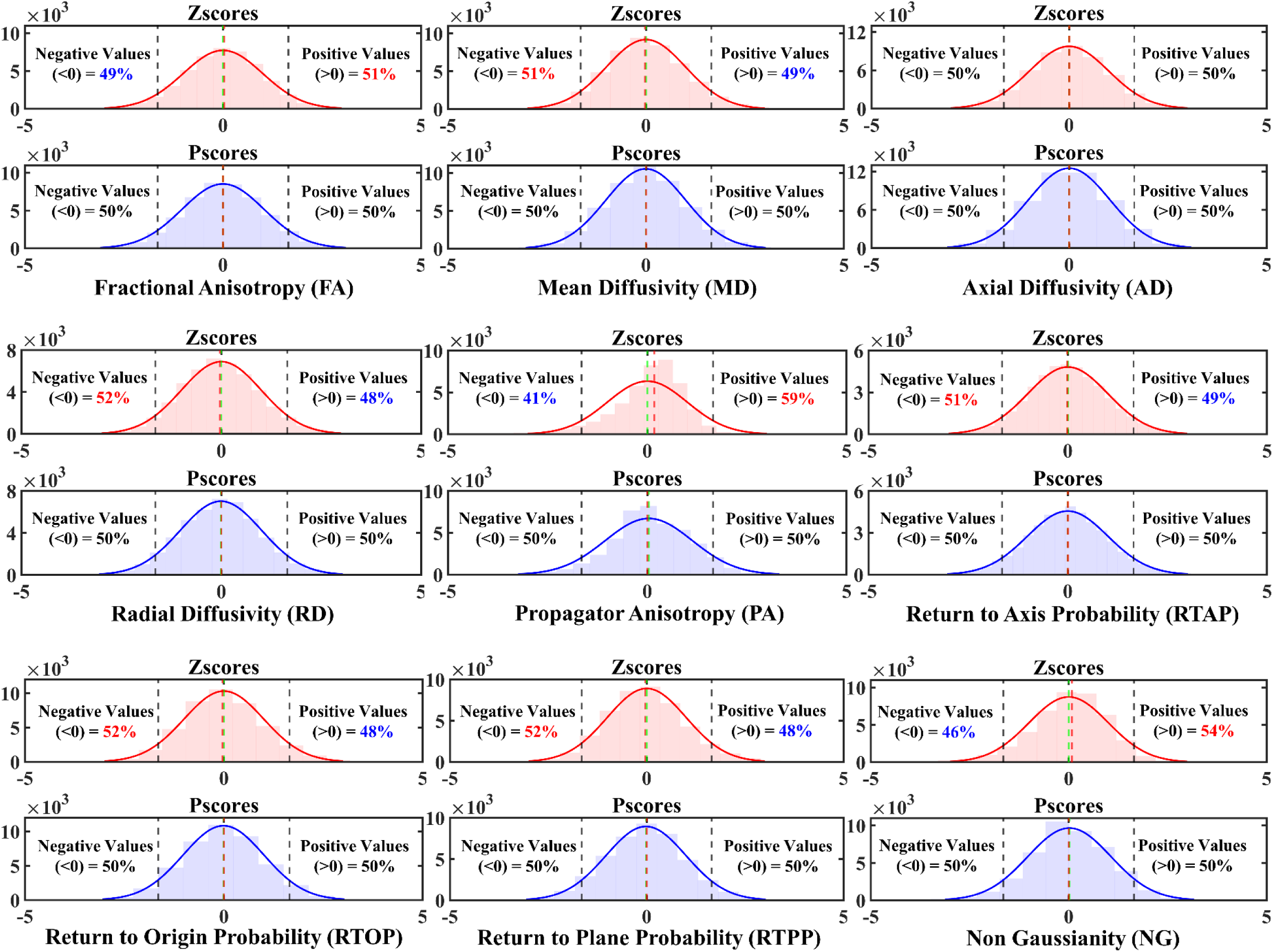
Comparing distributions of Zscores against Pscores across all WM ROIs per dMRI metric. For each dMRI metric, a pair of distributions are shown – one with Zscore (top, light red) histograms and one for Pscore (bottom, light blue) histograms. The values in these distributions came from concatenating all the scores across all ROIs into a single vector. It was done to show if there was an overall imbalance in the distribution from the entire WM. A normal density curve (solid red and blue) was also fit on top of each distribution, respectively, to assess which normalization method attained a closer fit to a Gaussian distribution. Except for AD, where the Zscore distribution was approximately Gaussian, an imbalance of negative and positive scores was observed for the distributions from all other dMRI metrics. The percentage of values above/below the 0-line are provided on either side of each distribution plot. An imbalance was indicated by a ‘> 50%’ and ‘< 50%’ value on either side of the 0-line. An increase above 50% is indicated in bold red and a decrease below 50% is indicated in bold blue numbers. The Zscore distribution of FA (top left, light red), for instance, showed an overall imbalance with 51% of the total observations above 0 and 49% below it. This means there were at least 460 (1% of 46080) more positive Zscores than negative Zscores in the distribution. The mean (green vertical dashed line) and the median (red vertical dashed line) also appeared misaligned. This pattern was also observed for PA and NG distributions. On the contrary, Zscores from MD, RD, RTAP, RTOP and RTPP showed an overall positive skew, with more negative (> 50%) and less positive (< 50%) values around the zero line. Pscores, however, consistently maintained an equal distribution of 50% negative and positive values on either side. The mean and median lines aligned, and the histograms attained a closer fit to the normal density curve (solid blue) compared to the Zscores (solid red).

### Assessing the Imbalance in Extreme Values at the Tails

Table 1 shows the imbalance of extreme values at the tail ends of the Zscore distributions and the correct balance of extreme values from the Pscore distributions. For each dMRI metric, it shows the percentage of extreme values above the 95^th^ (PEV_>95_(%), Z > 1.645) and below the 5^th^ (PEV_<5_(%), Z < −1.645) percentile boundaries for both these normalization methods. Except for AD, where the imbalance was negligible but still < 5%, Zscores from every other dMRI metric showed an imbalance of extreme values in the left or right tails. For example, FA, NG, and PA had progressively larger imbalance between negative and positive values leading to negative skews (Figure 3), and as a result, a subsequent increase in the percentage of negative extreme values. Compared to ‘PEV_>95_(%)’, the ‘PEV_<5_(%)’ values in these three metrics were higher by approximately 1.2 (4.9/4.1), 1.9 (5.4/2.8), and 5.8 (5.8/1) times, respectively. On the other hand, MD, RD, RTAP, RTOP, and RTAP showed positive skews for Zscores (Figure 3), leading to an increase in positive extreme values. All ‘PEV_>95_(%)’ values for these metrics were > 5%, while all ‘PEV_<5_(%)’ values were < 5%. Contrarily, for the same dMRI metrics, the ‘PEV_<5_(%)’ and ‘PEV_>95_(%)’ values for Pscores maintained a balanced 5% extreme values in both tails.

**Table 1.**
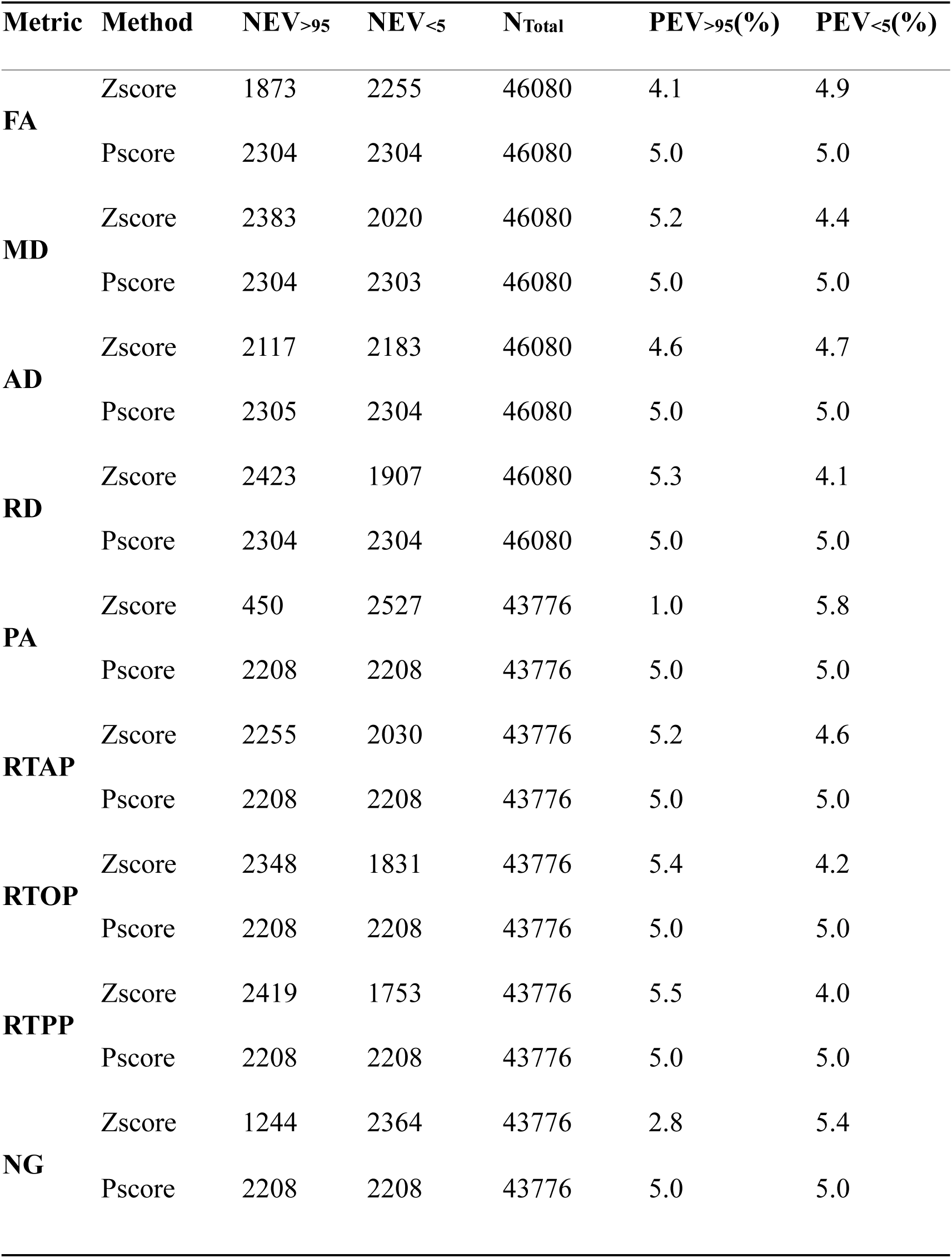
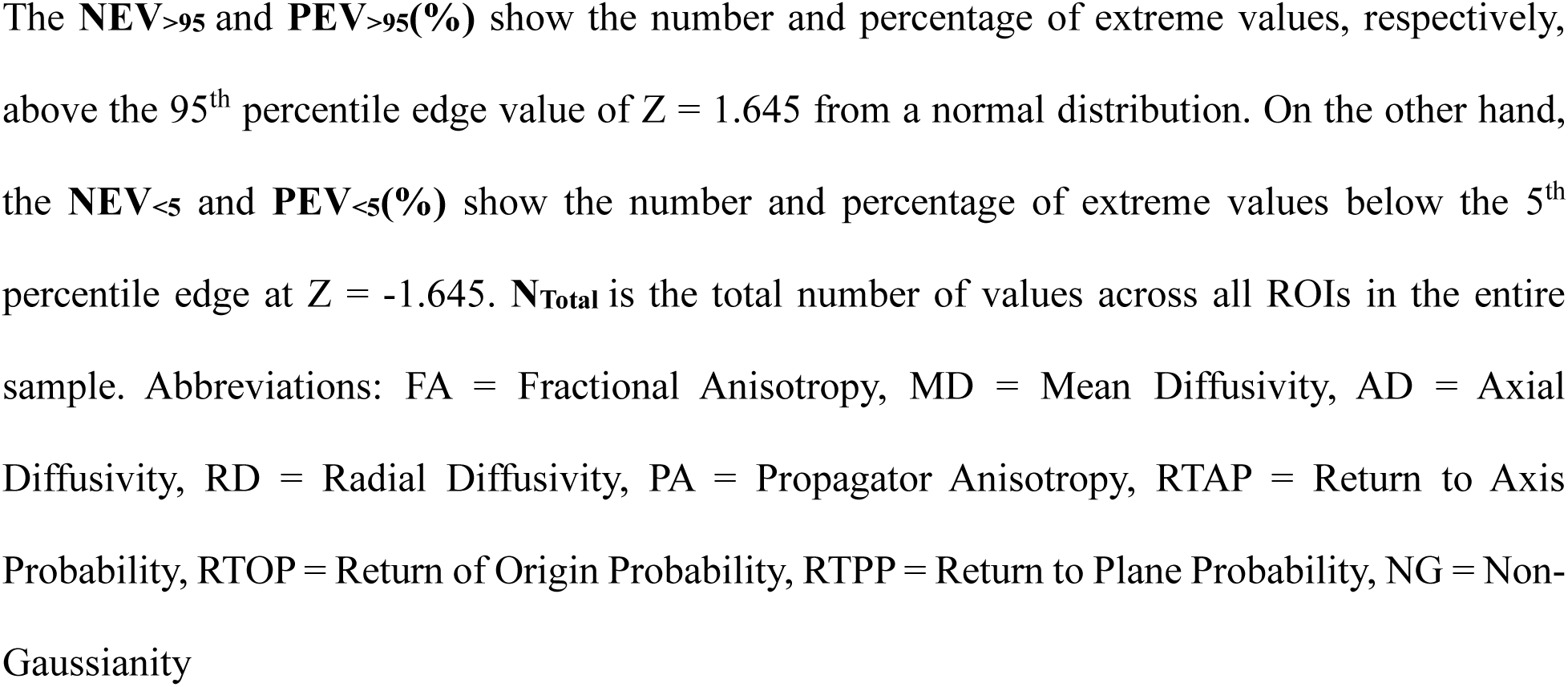
Assessing the Extreme Value Imbalance in Zscores Compared to Pscores.

### Bootstrapping the HCP Sample

Figure 4 shows a heatmap of PA generated from one iteration of a random sampling of 100 HCP participants. The heatmap demonstrates any systematic increase in extreme values in individuals (columns) across multiple ROIs. It also highlights any increase in extreme values present in an ROI (rows) across the normative sample. The heatmap of Zscores (top) shows a large number of negative extreme values and very few positive extreme values, whereas the Pscore heatmap (bottom) shows a proper balance of positive and negative extreme values, as expected of a normative sample. For simplicity, we showcase a heatmap comprising only 100 participants.

**Figure 4.**
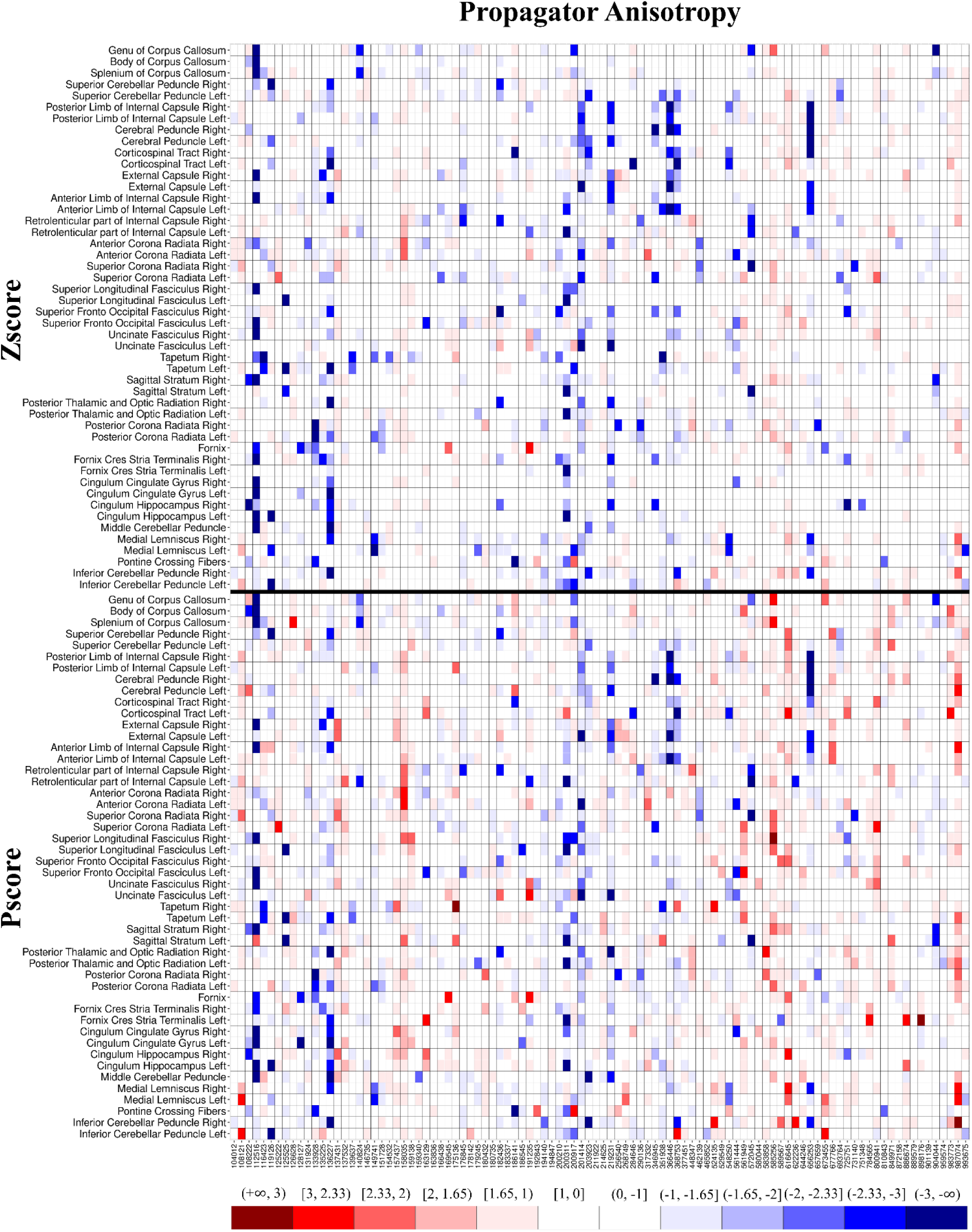
Heatmap comparing Zscores and Pscores from a single iteration of randomly selected sample of 100 participants for the Propagator Anisotropy. The horizontal black line separates the Zscores (top) from the Pscores (bottom). The columns represent individual participants, and the rows represent the regions of interest (ROIs). The x-axis shows the labels of the 100 participants randomly selected from the HCP. The range of Pscores are shown in the colorbar. The parenthesis ‘(’ in a colorbar tile means that the corresponding shade is exclusive of the value next to it, whereas the bracket ‘]’ means the value is inclusive. For example, the ‘[1.65,1)’ label on top of the lightest shade of red means it represents values in the range ‘1 < Z ≤ 1.65’ and so on for others. Darker shades of blue and red represented more extreme negative and positive values, respectively. An individual with a dark blue tile with the range label (−3, -∞) would correspond to Z < −3 from a normal distribution and vice-versa for a dark red shade. The large number of negative extreme values in Zscores (top) was quite conspicuous and a balanced positive and negative extreme values in the Pscores (bottom) was evident just from visual inspection.

Table 2 shows similar quantities as Table 1, except for PA, with 20 iterations of random sampling 100 HCP participants. The rationale to highlight PA was that it showed the highest imbalance in extreme values at the tails, among all dMRI metrics (Figure 3 and Table 1). For all 20 iterations, Zscores of PA showed large proportions of negative extreme values (all ‘PEV_<5_(%)’ > 5% and all ‘PEV_>95_(%)’ < 2%) with on average > 4.3 (6.1/1.4) times more negative extreme values than positives. Contrarily, Pscores robustly maintained 5% extreme values in both tails for all 20 iterations. Table S2 of the supplementary material provides a summary of the bootstrapping assessment of all dMRI metrics across 100 iterations.

**Table 2.**
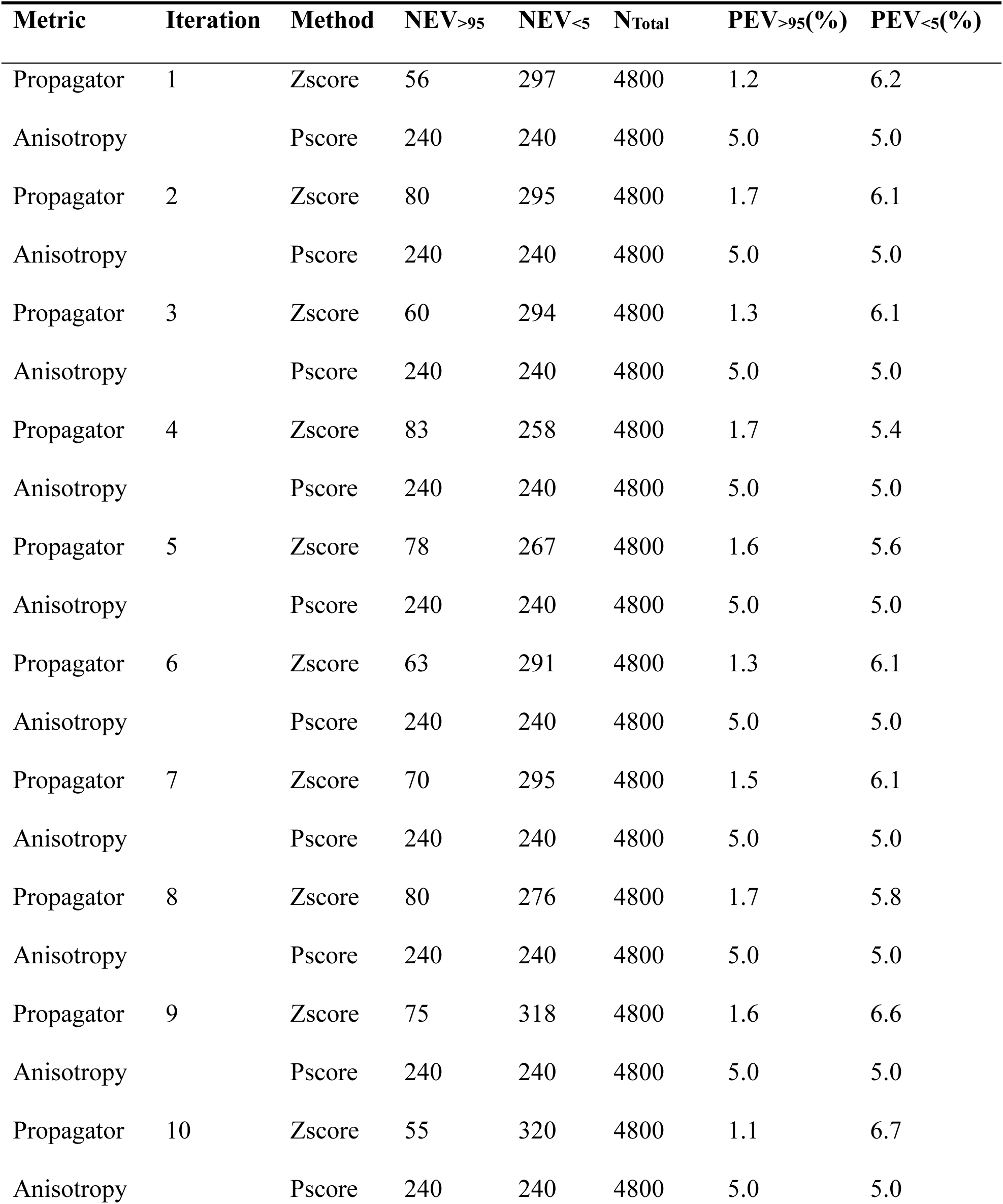
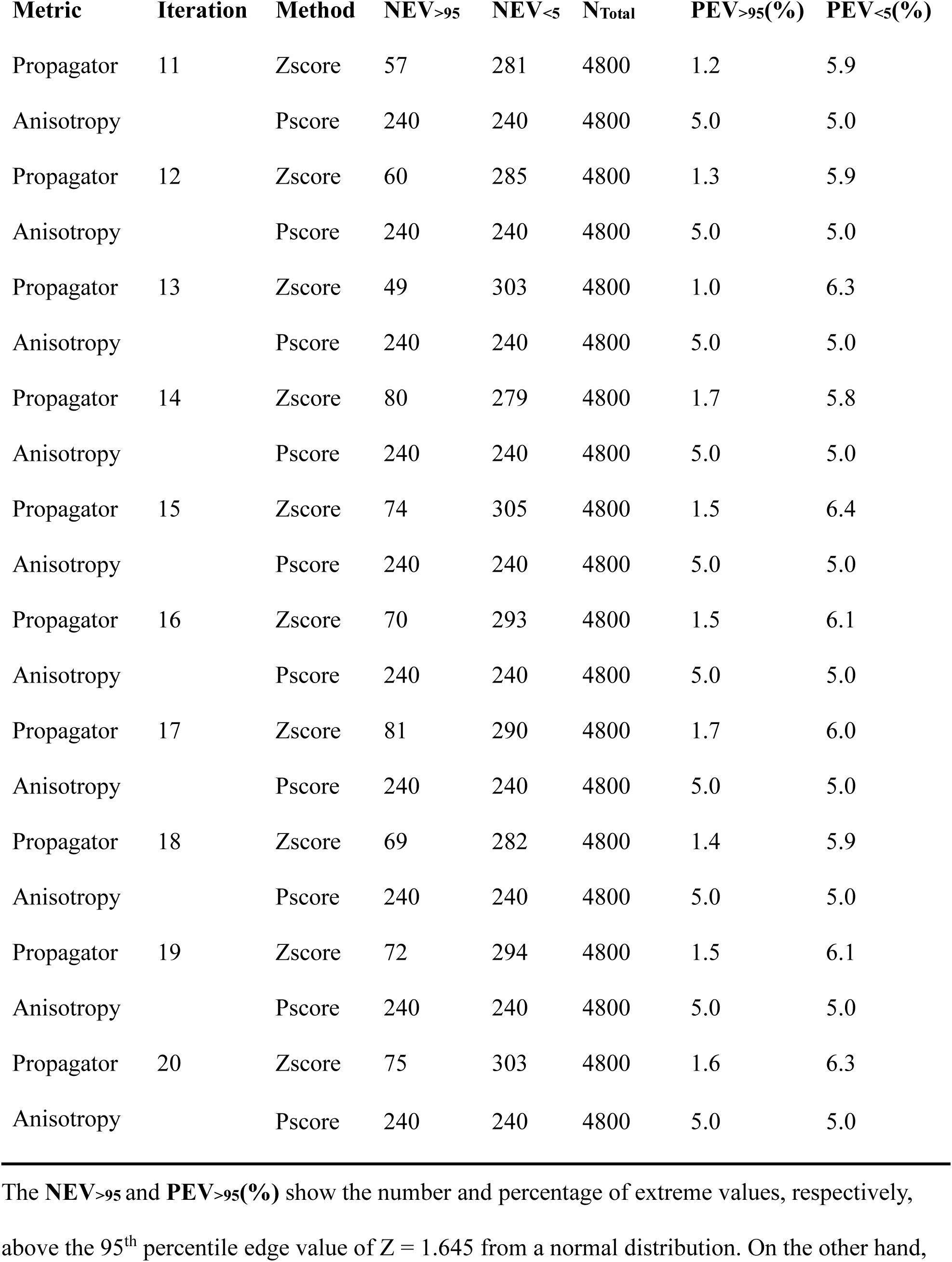
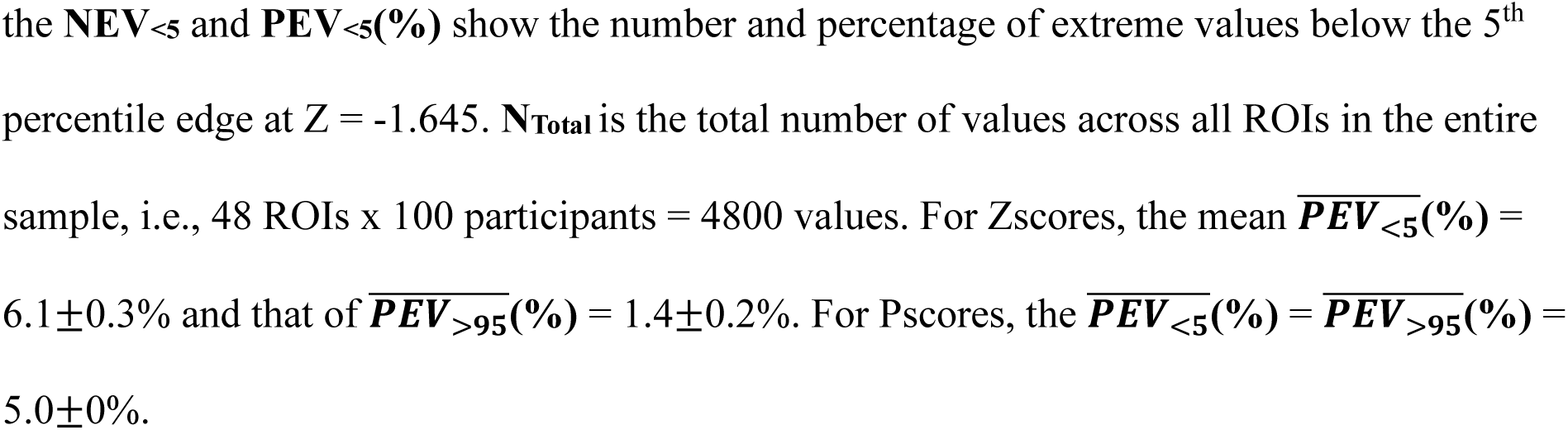
Comparing the Extreme Value Imbalance in Zscores vs. Pscores of Propagator Anisotropy for 20 Iterations of Bootstrapping 100 HCP Participants.

## DISCUSSION

Using the large-scale, high-resolution HCP dataset, we showed that the distributions of DT and MAP metrics derived from dMRI, tend to be non-Gaussian. We also showed that this may lead to an imbalance in the extreme values at the tails of a normative distribution, when Zscores are used. We proposed a novel percentile-based metric, the ‘Pscore’, which was less sensitive to these non-Gaussian distributions, both at the ROI level and for overall WM. We further documented the robustness of this method in smaller samples using bootstrapping. We replicated our previous findings from a pilot study and systematically validated this method using the HCP-YA cohort.

Our assessment on the metric distributions showed that diffusivity (e.g., MD, RD) tended to be positively skewed, whereas anisotropy (e.g., FA, PA) tended to be negatively skewed. It is difficult to unequivocally identify a primary source for these findings. However, given the narrow age range of the examined population, it is unlikely to be due to aging-related effects that have been invoked for large-scale public datasets (1,3,31). A possible explanation for these skewed distributions could be related to the presence of a small percentage of fast diffusing water molecules with isotropic diffusion behavior that have been reported in healthy brain parenchyma (32). This fast diffusing water compartment has the same diffusion signature as cerebrospinal fluid (CSF) partial volume contamination (32). Moreover, CSF contamination within a ROI can lead to higher MD (positive skew) because the diffusivity of CSF is at least four times higher than that of the brain parenchyma (33). On the other hand, the anisotropy of CSF is virtually 0, which would cause the anisotropy of the WM tracts to be lower (negative skew). These considerations suggest that skewed values may be inherently driven by underlying biological characteristics of the brain and not be necessarily related to heterogeneity in the demographics. One of our rationales for choosing the HCP young-adult sample was its well-balanced homogeneity in demographics (e.g., age and sex).

Our findings may be of particular interest to investigators assessing individual deviations against a skewed normative database. Normative models are typically built from very large samples and often incorporate Gaussian process models (1,3). This can be counterintuitive as scaling becomes a big challenge, as sample sizes get much bigger (3). Some methods have been proposed based on machine learning to address this issue, but with some caveats of pre-tuning and modeling complexity (34–36). Indeed, one can simply assess a quantity, e.g., an individual’s height, by comparing it against the percentiles generated from a sufficiently large population. However, aside from other confounds, neuroimaging studies are often limited by the sample size (median n = 23, according to (37)) and typically consist of < 100 participants. In a healthy cohort of 48 controls, we had previously shown that normative data generated from dMRI metrics deviate from normality and suffer the consequences of having unbalanced tail ends when Zscores are used (10). The study also underscored a key advantage of the ‘Pscore’ approach – its ability to reliably estimate individual deviations in small samples (10). Therefore, Pscores offer a practical solution to studies with more realistic sample goals (e.g., n = 50 – 150). It can help investigators to accurately assess individuals by building site-specific normative databases from neuroimaging data that do not conform to Gaussianity.

Deviation from Gaussianity leads to biases in the peripheral centiles of a normative distribution and inaccurate inferences (1). Zscores are commonly used for such inferences, and we have established the issues with extreme values that manifest if the issue of Gaussianity is not addressed. There are approaches that have been adopted to overcome non-Gaussian traits due to sample heterogeneity, implementing various modes of Bayesian linear regressions (1,38). The ‘Pscore’ approach addresses the issue of Gaussianity at the level of the data distribution itself. By leveraging the median as reference and using percentile ranks, it avoids the imbalance issue at the tails arising from the asymmetry in the distribution. Therefore, it can be an important addition to the various arsenal of contemporary normalization techniques (39). Pscores provide a simple platform to normalize raw data and scale them to standardized Zscores derived from a normal distribution. This is consequential because conforming to normally distributed Zscores facilitates statistical relevance to inferences made on individuals. It allows a meaningful interpretation of an individual’s position, especially when considering the extreme ends of a distribution.

When an individual, such as a patient, is expected to be at the extremities of a normative distribution, it can be considered as a rare event, as most individuals are not expected to be positioned there. Extreme value theorem and its subtypes can prove useful in such cases by fitting asymmetric curves on extreme value distributions derived from such rare events (40). This has been applied in neuroimaging applications to assess individuals from highly heterogeneous clinical cohorts using normative modeling (1,3). The process involves generating normative probability maps (NPMs) and computing a Zscore per brain region/voxel that normalizes the difference between the true normative and predicted individual response with their corresponding variances. These Zscores are then usually used to run univariate statistical tests with multiple comparison adjustments. Our expectation is that Pscores can prove very useful in these applications and provide more accurate estimations for extreme events compared to Zscores.

### Limitations

The Pscore computation relies on percentile ranks of every participant in the normative sample. Therefore, the sample size may be a limiting factor on the confidence and precision of the percentile edges. For example, in our pilot analysis of 48 controls (10), the smallest resolution of each percentile was 2.083 or ∼ 2% (100/48). Since the 5^th^ and 95^th^ percentile edges were used, there was some statistical uncertainty to the exact values representing these percentiles. However, the Pscores still maintained a balanced number of extreme values in the two tails. This issue obviously recedes as sample sizes get bigger, as we see in the current context (n = 961), where the percentile resolutions were much finer (∼ 0.1%) and allowed very precise edge measurements. Furthermore, the limitation can be overcome even with smaller samples if precise and even percentile edge measurements can be performed. Regarding the bootstrapping step for instance, the percentile resolutions were exactly even at 1% (100/100) and Pscores robustly maintained 5% extreme values over 100 iterations for all dMRI metrics.

Another limitation for Pscores is the selection of the percentile edges. We used the 5^th^ and 95^th^ percentile edges because they are statistically relevant and allow at least 10% of extreme values in the whole distribution. Choice of a more extreme percentile edge may affect the normalization accuracy. An appropriate alternative could be to look at the area under the curve, instead of point values at percentile edges. That way the computation will be done on a continuous scale, and it also preserves the feature of Pscores to address the issue at the level of the distribution, without making any prior assumption of Gaussianity. This is part of our future project using various samples of multimodal neuroimaging data.

### Conclusion

Although the ‘Pscore’ approach was tested on dMRI metrics, the method itself is not data selective. The ‘Pscore’ method, instead looks at the data by virtue of its distribution. Pscores can control the imbalance arising from the non-Gaussianity in the distribution and more accurately estimate individual deviations from a normative database. The robustness of Pscores observed for small samples in this study indicates its potential application to assess individuals in studies with smaller databases. The application of this method on data from clinical and other modalities, such as structural volumetric and functional MRI, may warrant future investigation because metrics from these paradigms also suffer from small sample limitations and non-Gaussian distributions.

## Acknowledgments

Data were provided [in part] by the Human Connectome Project, WU-Minn Consortium (Principal Investigators: David Van Essen and Kamil Ugurbil; 1U54MH091657) funded by the 16 NIH Institutes and Centers that support the NIH Blueprint for Neuroscience Research; and by the McDonnell Center for Systems Neuroscience at Washington University.

