## Supplementary Materials for "‘Pscore’ - A Novel Percentile-Based Metric to Accurately Assess Individual Deviations in Non-Gaussian Distributions of Quantitative MRI Metrics"

**Table S1. List of Regions of Interest (ROIs) Used in the Current Study**

| ROI Numbers | ROI Names | Number of Voxels | Volume of ROI  /mm^3^ |
| --- | --- | --- | --- |
| 1 | Middle Cerebellar Peduncle | 2320 | 2320 |
| 2 | Pontine Crossing Fibers | 480 | 480 |
| 3 | Genu of Corpus Callosum | 1285 | 1285 |
| 4 | Body of Corpus Callosum | 3283 | 3283 |
| 5 | Splenium of Corpus Callosum | 3432 | 3432 |
| 6 | Fornix | 87 | 87 |
| 7 | Corticospinal Tract Right | 344 | 344 |
| 8 | Corticospinal Tract Left | 253 | 253 |
| 9 | Medial Lemniscus Right | 188 | 188 |
| 10 | Medial Lemniscus Left | 163 | 163 |
| 11 | Inferior Cerebellar Peduncle Right | 428 | 428 |
| 12 | Inferior Cerebellar Peduncle Left | 372 | 372 |
| 13 | Superior Cerebellar Peduncle Right | 179 | 179 |
| 14 | Superior Cerebellar Peduncle Left | 166 | 166 |
| 15 | Right Cerebral Peduncle | 333 | 333 |
| 16 | Left Cerebral Peduncle | 317 | 317 |
| 17 | Anterior Limb of Internal Capsule Right | 490 | 490 |
| 18 | Anterior Limb of Internal Capsule Left | 572 | 572 |
| 19 | Posterior Limb of Internal Capsule Right | 832 | 832 |
| 20 | Posterior Limb of Internal Capsule Left | 624 | 624 |
| 21 | Retrolenticular part of Internal Capsule Right | 614 | 614 |
| 22 | Retrolenticular part of Internal Capsule Left | 466 | 466 |
| 23 | Anterior Corona Radiata Right | 1529 | 1529 |
| 24 | Anterior Corona Radiata Left | 1319 | 1319 |
| 25 | Superior Corona Radiata Right | 1934 | 1934 |
| 26 | Superior Corona Radiata Left | 1343 | 1343 |
| 27 | Posterior Corona Radiata Right | 586 | 586 |
| 28 | Posterior Corona Radiata Left | 630 | 630 |
| 29 | Posterior Thalamic and Optic Radiation Right | 660 | 660 |
| 30 | Posterior Thalamic and Optic Radiation Left | 806 | 806 |
| 31 | Sagittal Stratum Right | 938 | 938 |
| 32 | Sagittal Stratum Left | 602 | 602 |
| 33 | External Capsule Right | 1461 | 1461 |
| 34 | External Capsule Left | 921 | 921 |
| 35 | Cingulum Cingulate Gyrus Right | 758 | 758 |
| 36 | Cingulum Cingulate Gyrus Left | 649 | 649 |
| 37 | Cingulum Hippocampus Right | 324 | 324 |
| 38 | Cingulum Hippocampus Left | 261 | 261 |
| 39 | Fornix Cres Stria Terminalis Right | 184 | 184 |
| 40 | Fornix Cres Stria Terminalis Left | 192 | 192 |
| ROI Numbers | **ROI Names** | **Number of Voxels** | **Volume of ROI**  **/mm^3^** |
| 41 | Superior Longitudinal Fasciculus Right | 1047 | 1047 |
| 42 | Superior Longitudinal Fasciculus Left | 886 | 886 |
| 43 | Superior Fronto Occipital Fasciculus Right | 100 | 100 |
| 44 | Superior Fronto Occipital Fasciculus Left | 80 | 80 |
| 45 | Uncinate Fasciculus Right | 123 | 123 |
| 46 | Uncinate Fasciculus Left | 83 | 83 |
| 47 | Tapetum Right | 74 | 74 |
| 48 | Tapetum Left | 86 | 86 |

The **‘Number of Voxels’** represents the total sum of the number of voxels within an ROI. The **‘Volume of ROI’** represents the total volume in mm^3^ of each ROI. The values in the two columns appear identical because the volumes were computed on an isotropic 1 mm resolution, giving each voxel a unit volume of 1 mm^3^.

**Table S2. Assessing Extreme Value Imbalance in Zscores vs Pscores of all Diffusion MRI Metrics from Bootstrapping Across 100 Iterations.**

| **Metric** | **Method** | $\bar{\mathbf{NEV}_{\mathbf{>}\mathbf{95}}}$ | $\bar{\mathbf{NEV}_{\mathbf{<}\mathbf{5}}}$ | $\bar{\mathbf{N}_{\boldsymbol{Total}}}$ | $\bar{\boldsymbol{PEV}_{\boldsymbol{>}\boldsymbol{95}}}$**(%)**  Mean ± SD | $\bar{\boldsymbol{PEV}_{\boldsymbol{<}\boldsymbol{5}}}$**(%)**  Mean ± SD |
| --- | --- | --- | --- | --- | --- | --- |
| Fractional Anisotropy (FA) | Zscore | 206 | 246 | 4800 | 4.3 ± 0.4 | 5.1 ± 0.3 |
|  | Pscore | 240 | 240 | 4800 | 5.0 ± 0.0 | 5.0 ± 0.0 |
| Mean  Diffusivity (MD) | Zscore | 254 | 219 | 4800 | 5.3 ± 0.3 | 4.6 ± 0.3 |
|  | Pscore | 240 | 240 | 4800 | 5.0 ± 0.0 | 5.0 ± 0.0 |
| Axial  Diffusivity (AD) | Zscore | 228 | 238 | 4800 | 4.8 ± 0.3 | 5.0 ± 0.3 |
|  | Pscore | 240 | 240 | 4800 | 5.0 ± 0.0 | 5.0 ± 0.0 |
| Radial  Diffusivity (RD) | Zscore | 256 | 206 | 4800 | 5.3 ± 0.3 | 4.3 ± 0.3 |
|  | Pscore | 240 | 240 | 4800 | 5.0 ± 0.0 | 5.0 ± 0.0 |
| Propagator Anisotropy (PA) | Zscore | 65 | 289 | 4800 | 1.4 ± 0.2 | 6.0 ± 0.3 |
|  | Pscore | 240 | 240 | 4800 | 5.0 ± 0.0 | 5.0 ± 0.0 |
| Return to Axis Probability (RTAP) | Zscore | 252 | 226 | 4800 | 5.3 ± 0.3 | 4.7 ± 0.3 |
|  | Pscore | 240 | 240 | 4800 | 5.0 ± 0.0 | 5.0 ± 0.0 |
| Return to Origin Probability (RTOP) | Zscore | 266 | 211 | 4800 | 5.5 ± 0.3 | 4.4 ± 0.3 |
|  | Pscore | 240 | 240 | 4800 | 5.0 ± 0.0 | 5.0 ± 0.0 |
| Return to Plane Probability (RTPP) | Zscore | 268 | 198 | 4800 | 5.6 ± 0.2 | 4.1 ± 0.3 |
|  | Pscore | 240 | 240 | 4800 | 5.0 ± 0.0 | 5.0 ± 0.0 |
| Non  Gaussianity (NG) | Zscore | 151 | 271 | 4800 | 3.1 ± 0.3 | 5.6 ± 0.2 |
|  | Pscore | 240 | 240 | 4800 | 5.0 ± 0.0 | 5.0 ± 0.0 |

The $\bar{\mathbf{NEV}_{\mathbf{>95}}}$ and $\bar{\boldsymbol{PEV}_{\boldsymbol{>95}}}$ show the mean number and percentage of extreme values, respectively, above the 95^th^ percentile edge value of Z = 1.645 from a normal distribution. The ‘accent’ bar on top represents the ‘mean’ across 100 iterations. Similarly, the $\bar{\mathbf{NEV}_{\mathbf{<5}}}$ and $\bar{\boldsymbol{PEV}_{\boldsymbol{<5}}}$ show the mean number and percentage of extreme values below the 5^th^ percentile edge value of Z = -1.645. $\bar{\mathbf{N}_{\boldsymbol{Total}}}$ is the mean across 100 iterations for the total number of values across all ROIs in the randomly selected sample of 100 participants. For the $\bar{\boldsymbol{PEV}_{\boldsymbol{>95}}}$ and $\bar{\boldsymbol{PEV}_{\boldsymbol{<5}}}$ values**,** both the mean and standard deviation (SD) across 100 iterations, are shown for each method.

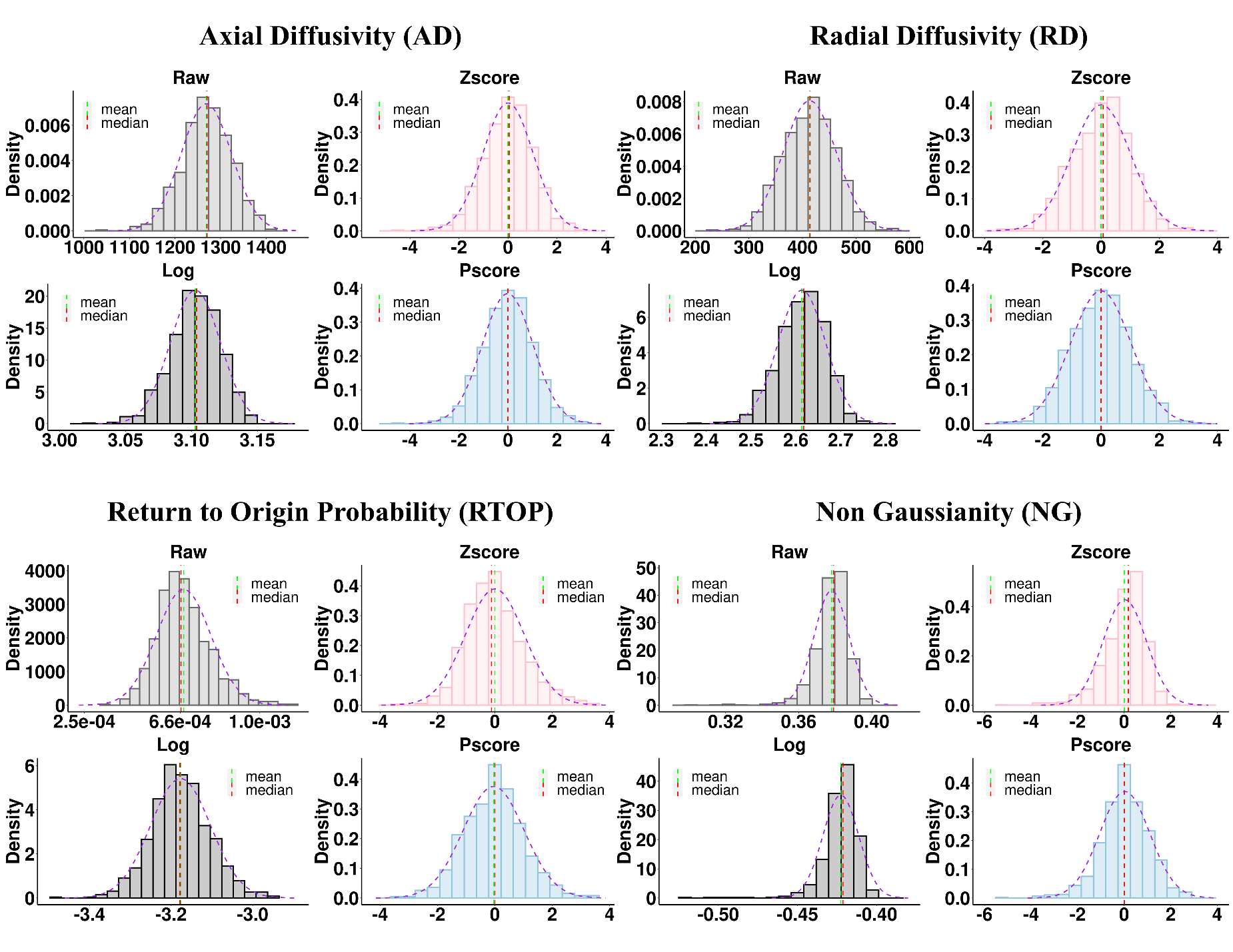
**Figure S1. Comparing distributions across different normalization methods for two DT and two MAP metrics from a representative ROI: Anterior Limb of Internal Capsule Right.** The DT metrics are shown in the top row: AD (left) and RD (right); and the MAP metrics are on the bottom row: RTOP (left) and NG (right). For each metric, four distributions are shown. The ‘Raw’ (light gray) distribution panel represents average metric values. The ‘Log’ (dark gray), ‘Zscore’ (light red) and ‘Pscore’ (light blue) distribution panels represent the log transformed, standardized Zscores and the proposed metric: Pscores, respectively. AD (top left), RD (top right) and NG (bottom right) were negatively skewed, with NG showing the strongest skew. The green and red dashed lines represent the mean and median of the distributions. For AD, RD and NG, the ‘mean’ underestimated the most common values and appeared before the median and did not coincide well with the mode of the distribution for ‘Raw’, ‘Log’ and ‘Zscore’ panels. However, the ‘Pscore’ panel shows that all three measures of central tendencies aligned well and attained a closer fit to the normal density curve (dashed purple). On the other hand, RTOP (bottom left) was positively skewed and the ‘mean’ overestimated the most common values for ‘Raw’, ‘Log’ and ‘Zscore’ panels, but the ‘Pscore’ panel showed a closer alignment of mean, mode and median and a better fit to the normal density curve.

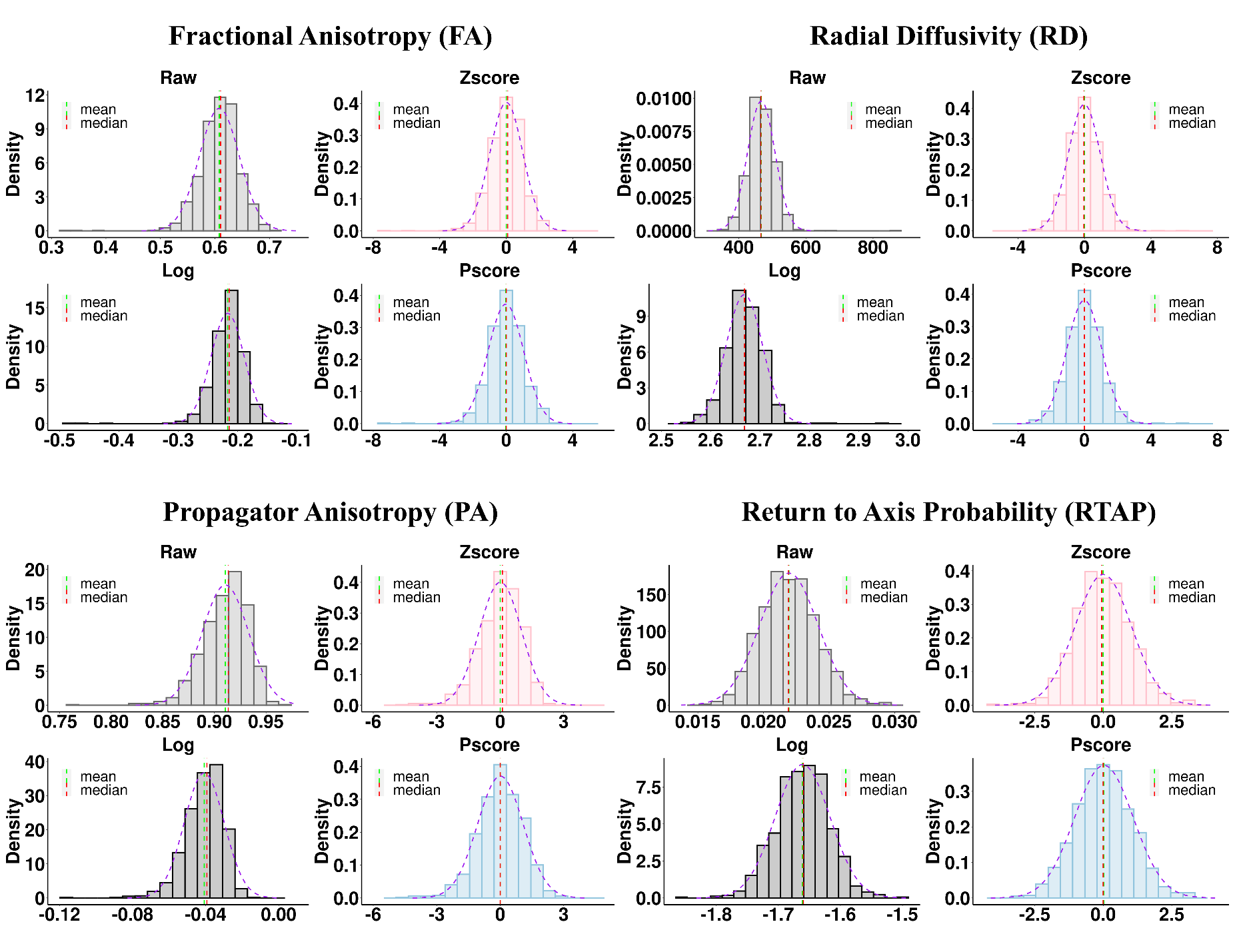
**Figure S2. Comparing distributions across different normalization methods for two DT and two MAP metrics from a representative ROI: Medial Lemniscus Left.** Similar descriptions as Figure S1 applies except for DT metrics – FA (top left) and RD (top right); and MAP metrics – PA (bottom left) and RTAP (bottom right). FA and PA were negatively skewed. For these metrics, the ‘mean’ underestimated the most common values and appeared before the median and the mode of the distribution for ‘Raw’, ‘Log’ and ‘Zscore’ panels. However, the ‘Pscore’ panel shows that all three moments aligned well and attained a closer fit to the normal density curve (dashed purple). On the other hand, RD and RTAP were positively skewed and the ‘mean’ overestimated the most common values for ‘Raw’, ‘Log’ and ‘Zscore’ panels. However, the Pscores showed consistent alignment of the mean, mode and median and a closer fit to the normal density curve.
